# Measuring adaptation dynamics to hydrogen peroxide in single human cells using fluorescent reporters

**DOI:** 10.1101/2020.11.14.382911

**Authors:** Dana Simiuc, Fatima Dahmani, Alexandra Pruvost, Marie Guilbert, Mathilde Brulé, Chann Lagadec, Quentin Thommen, Benjamin Pfeuty, Emmanuel Courtade, François Anquez

## Abstract

We developed an experimental methodology to monitor response dynamics of single human cells to hydrogen peroxide. Our approach is based on fluidic control of both magnitude and time-evolution of the external perturbation, and on high-throughput imaging of intracellular fluorescent redox reporters. We applied step stimulus to MCF7 cells with hydrogen peroxide concentration in the range of 10 to 500*μM*. First, our data highlights dynamic adaptation of Reactive Oxygen Species (ROS) scavenging system at several time scales. Nicotinamide Adenine Dinucleotide Phosphate (NADPH) level is rapidly restored within 10 min after a transient decrease, while glutathione (GSH) redox potential is slowly driven back toward pre-stimulus level (within one hour). Extra-cellular glucose is necessary for adaptation of both NADPH level and GSH system. Second, our results also reveal large cell-to-cell variability in the dynamic response to external ROS. Our experimental approach is readily usable to monitor other cellular redox systems such as thioredoxins. As response-heterogeneity can lead to fractional killing, we finally discuss how our methodology can be an opportunity to link dynamics of ROS scavenging and cellular fate such as cell death.

## INTRODUCTION

Exposure to an external perturbation, such as heat or radiation, can cause damages to cellular macro-molecules, which can in turn lead to cell death. Such a perturbation is called stress, and human cells have inherited the ability to respond to stress. They do so by triggering a biochemical activity that prevents and/or repairs damages. Stress and associated cellular response are involved in the process of aging [1], and can be responsible for carcinogenesis [2, 3] and degenerative diseases [4]. Among all types of injuries, oxidative stress that is due to excess of Reactive Oxygen Species (ROS) is of particular interest. Indeed, ROS are known to contribute to the efficiency of radiotherapy [5] and chemotherapy [6, 7], and photodynamic therapy is a treatment modality that uses light to specifically produce ROS [8]. Oxidative stress response is complex and involves together several pathways such as activation transcriptional programs [9, 10], post-transcriptional modifications [11, 12], *de novo* synthesis of antioxidants [13] and rerouting of metabolic fluxes to counter the redox imbalance [14, 15].

At the front line against the excess of ROS are scavenging systems that involve enzymatic and non-enzymatic antioxidants [16, 17]. Among these, the glutathione system plays a central role. It uses glutathione (GSH) as a reducing cofactor for enzymes to scavenge ROS and detoxify oxidized proteins [16, 17]. The glutathione system is powered by NADPH, which is itself renewed by glucose metabolism *via* several routes including the Pentose Phosphate Pathway (PPP) [18, 16]. In recent years several works have reported a highly dynamic response to oxidative load, involving several time scales: rapid increase of flux in the oxidative branch of the Pentose Phosphate Pathway (PPP) [14, 19] and slower metabolic rerouting toward the PPP [15, 20, 21, 22], and even slower transcriptional responses [23, 24]. All these data reveal fine regulations that can lead to adaptation to the perturbation: after a transient response some quantities of the system tends to come back close to prestimulus state [25, 23, 15, 14, 19]. Stress response is thus a highly dynamic process, and the understanding of regulations, including their dynamics, may help to improve therapeutic strategies that, for example, use drugs to sensitize cancer cells to treatment[26].

Adaptation to an external stimulus are common features of biological systems [27, 28, 17, 29]. A limited amount of biochemical network schemes allow such an adaptation [30]. Temporal control of the stimulus together with time-resolved monitoring of the response have been successfully employed to decipher this types of regulatory networks [31, 32, 33, 34, 35]. In this work, we used hydrogen peroxide (*H*_2_*O*_2_) as a model perturbation for oxidative stress. In cells *H*_2_*O*_2_ reacts with thiols and, in particular, cysteine residues of proteins [36]. It also induces DNA damages, including single strand breaks, *via* Fenton chemistry [37, 38]. *H*_2_*O*_2_ is scavenged by catalase, the thioredoxin system, and the glutathione system [17]. We have developed a protocol, based on fluidics, to control both amplitude and time-evolution of *H*_2_*O*_2_ stimuli. We applied step stimulus of *H*_2_*O*_2_ to breast cancer cells MCF7. Abundance of proteins involved in metabolic regulation and ROS scavenging is known to be very different in the cancerous cell line MCF7 compared to its non-cancerous equivalent MCF10A [39]. And we used fluorescent reporters and optical microscopy to monitor dynamics of cellular response following stimulation. On the one hand, we used GRx1-roGFP2 that senses glutathione redox potential [40], and revealed a slowly adapting phenotype upon prolonged exposure to *H_2_O_2_* (within one hour). After ruling out potential side effects, we showed that adaptation is dose-dependent. On the other hand, we monitored intracellular NADPH dynamics *via* its autofluorescence [41, 42]. NADPH exhibited faster adaptation (within minutes). Glucose-deprivation showed that the regulation observed for GSH is dependent of glucose uptake. Moreover, while NADPH-adaptation is not inhibited in the absence of glucose, we observed that the lack of adaptation of GSH coincides with a lower NADPH level at steady state in the absence of glucose. We finally discussed plausible mechanisms for the observed adaptation and the advantages of our methodology to monitor dynamics of oxidative stress response.

## RESULTS

### Control of external *H*_2_*O*_2_ stimulus reveals a potential regulation on ROS scavenging

In order to be able to highlight a potential adaptation, one needs to ensure a steady stimulus. *H*_2_*O*_2_ degradation in culture medium (see Figure SI 1A and reference [43]) renders this task more complicated. We indeed found a *H*_2_*O*_2_ half-life of ~ *1h* in extracellular medium and in the presence of cells. To counteract such a concentration decrease, our strategy was to renew extracellular *H*_2_*O*_2_ faster than it is degraded. For this purpose, we designed a fluidic system that allows us to expose cells to *H*_2_*O*_2_-free medium or to medium with a given *H*_2_*O*_2_ concentration (Figure 1A). Similar strategy, inspired from works on signal transduction in bacterial chemotaxis [27, 28], was applied by others to study stress response in yeast cells [23]. At a flow rate of 0.5*mL*/*min*, the medium in our chamber was fully renewed within one minute. This time scale is much faster than *H*_2_*O*_2_ half life in the presence of cells. Although even faster time scales can be reached with our system, we chose to limit the flow rate to 0.5*mL*/*min*. Doing so, shear stress does not induce signal transduction [44, 45]. Using propidium iodide (PI) staining, we verified that cells can survive for at least 24h in complete DMEM medium in our fluidic chamber. Our system enables good control on the temporal shape of the stimulus. At 0.5*mL*/*min* flow rate, we could produce a step stimulus with rise and decay time of ~ 1*min* (see Figure 1B).

**Figure 1:**
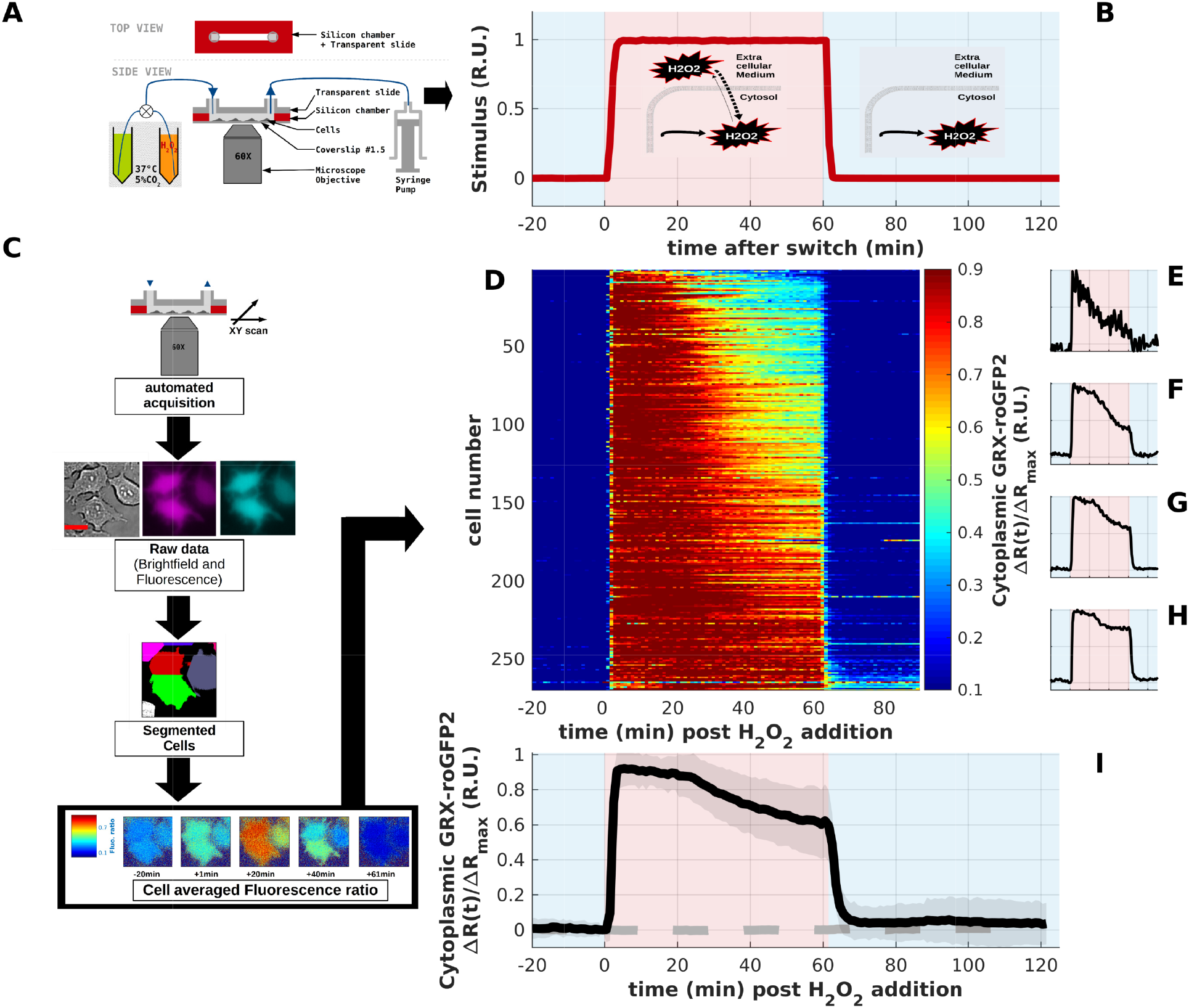
Controlling *H*_2_*O*_2_ stimulus and monitoring intracellular glutathione redox potential revealed a potential adaptation of ROS scavenging system: **A** - Schematic representation of our experimental system - The flow chamber was built by assembling a glass coverslip with a transparent slide *via* a silicon spacer. Adherent cells were seeded on the glass coverslip. From one side of the chamber, medium was pulled with a syringe pump at 0.5*mL*/*min*. A switch was connected on the other side of the chamber. It allowed to alternatively expose cells to *H*_2_*O*_2_ free medium or to a given *H*_2_*O*_2_ concentration. This system allowed to create a step stimulus (as plotted on panel B). **B** - The effective intracellular *H*_2_*O*_2_ production rate can be controlled by modulating external *H*_2_*O*_2_ concentration - Kinetics in our flow chamber was simulated by replacing *H*_2_*O*_2_ with a fluorescent dye (rhodamin 110). External medium concentration is plotted as a function of time. In the absence of external stimulus *H*_2_*O*_2_ is produced by cellular metabolism at rate Γ_*bas*_. In the presence of external *H*_2_*O*_2_ the total effective production rate is increased: Γ_*tot*_ = Γ_*bas*_ + *K_in_*.[*H*_2_*O*_2_]_*ext*_. **C** - Monitoring intracellular redox dynamics in single-cells - The chamber was placed on a custom-built automated microscope [51]. We imaged cells expressing various endogenous fluorescent redox probes such as Grx1-roGFP2, a sensor of intracellular glutathione redox potential [40]. By means of image processing and cell segmentation we produced single-cell time series of the quantity of interest. For example, we extracted kinetics of Grx1-roGFP2 fluorescence ratio, *R*, which reflects intracellular *H*_2_*O*_2_ concentration. **D-I** - *H*_2_*O*_2_ stimulation of MCF7 cells revealed dynamic adaptation - MCF7 cells expressing cytoplasmic Grx1-roGFP2 were imaged in the flow chamber. Cells were exposed to *H*_2_*O*_2_ free DPBS supplemented with 4.5g/L glucose for 30*min* then exposed to 100*μM H*_2_*O*_2_ in DPBS with 4.5g/L glucose for 60*min* and recovered in *H*_2_*O*_2_ free DPBS supplemented with 4.5g/L glucose for another 60*min*. We imaged 270 cells at 1 frame per *min*. We plotted change in Grx1-roGFP2 fluorescence ratio compared to its pre-stimulus level, Δ*R*, relative to maximum change upon *H*_2_*O*_2_ addition Δ*R_max_*. D - false color image displaying Δ*R*(*t*) / Δ*R_add_* as a function of time for all cells. E-H - Time traces of single-cells: #1, #66, #131 and #196. I - Change in Grx1-roGFP2 signal averaged over the whole cell population for stimulated cells (black line) and for a control experiment in which *H*_2_*O*_2_ was replaced by mq-water (gray dashed line). Grey shaded area represents standard deviation.

Another important aspect, if one wants to highlight a potential adaptation, is purity of the stimulus. As *H*_2_*O*_2_ reacts with chemical components of the culture medium (Figure SI 1A), one could expect generation of secondary species including reactive ones. To avoid such a side effect, we chose to work in DPBS supplemented with 4.5*g*/*L* glucose. We did not detect *H*_2_*O*_2_ degradation in DPBS and nor in the presence of glucose (see Figure SI 1C). Using our fluidic system, we could control the effective *H*_2_*O*_2_ intracellular production rate Γ. In the absence of extracellular *H*_2_*O*_2_, the basal production rate, Γ_*bas*_, is low with *H*_2_*O*_2_ predominantly produced as a side product of cell metabolism [46]. When stimulus is applied, penetration of extracellular *H*_2_*O*_2_ inside cells mimics an increase of an apparent *H*_2_*O*_2_ production rate. Effective intracellular production rate reaches a value Γ_*tot*_ > Γ_*bas*_ (see Figure 1B)[47, 48].

We have used fluorescence microscopy to monitor dynamics of endogenous redox processes [49]. In this work we have used: Grx1-roGFP2 reflecting glutathione (GSH/GSSG) redox potential [40] and NAD(P)H auto-fluorescence to monitor its intracellular concentration [50]. By scanning the sample with a motorized XY-stage our setup allowed automated acquisition of fluorescence images for several hundreds of cells over time [51]. Using image processing and analysis, we could retrieve redox dynamics for up to 1000 individual cells at 1 frame per minute (see Figure 1C).

We first applied a 1*h* - 100*μM H*_2_*O*_2_ step stimulus to MCF7 cells stably expressing cytosolic Grx1-roGFP2. We monitored Grx1-roGFP2 fluorescence at 520*nm* under excitation at 420*nm* and 482*nm*. The quantity of interest is the fluorescence ratio, *R*, (420*nm* versus 482*nm*) which increases when oxidation of Grx1-roGFP2 is favored. Experiments were carried in the presence of extracellular glucose. Control experiments, in which *H*_2_*O*_2_ was replaced by mq-water, did not show any change in *R* (gray dashed line in Figure 1I). Single-cell time series (Figure 1D-H and SI Video 1) showed that *R* increased shortly following stimulus addition because the GSH oxidation rate, *V_ox_*, increased. As expected, *R* rapidly reached a maximum when *V_ox_* equaled GSSG reduction rate, *V_red_*, ~ 5*min* after stimulus addition. More interestingly, we found that *R* slowly decreased after it reached a maximum even if extracellular *H*_2_*O*_2_ (and thus Γ_*tot*_) was kept constant. At 55*min* post stimulation *R* was, on average, diminished by ~ 40% of its maximum (Figure 1I). Step stimuli (1*h* - 100*μM H*_2_*O*_2_) were repeated three times. For each repeat, we found a diminution of *R* by ~ 40% of its maximum on average. The fact that *R* decreased after prolonged 100*μM H*_2_*O*_2_ exposure suggests that GSH redox potential tends to be restored during prolonged stimulation. Our data thus highlight a potential adaptation of ROS scavenging.

### pH calibration and analysis of Grx1-roGFP2 response confirm the existence of regulation and adaptative phenotype

At this stage, one must ensure that Grx1-roGFP2 signal, *R*, truly reflects glutathione redox potential, and that no other side effect can lead to misinterpretation of the data. We thus verified that cellular response does not depend on cell position in the chamber. We indeed did not find correlation between parameters characterizing *R* dynamics (defined later in the text) and position in the chamber. Moreover, using PI staining, we found that cell death does not arise before 4h post *H*_2_*O*_2_ stimulation in our experimental conditions, while the data presented here were recorded in not more than 2.5 hours post *H*_2_*O*_2_ exposure.

Cytosolic pH was shown to drop during *H*_2_*O*_2_ exposure [52]. Although Grx1-roGFP2 ratio, *R*, is pH-insensitive close to physiological conditions [40], *R* could be pH-dependent in extreme cases, because the spectral properties of the oxidized and reduced forms of the fluorophore (roGFP2) have different pH dependency [53]. We thus performed experiments to (*i*) estimate pH changes during stimulation, and to (*ii*) calibrate corresponding variations of Grx1-roGFP2 fluorescence ratio (see Supporting Information). First, using SypHer, a fluorescent pH-probe [54, 55], we measured pH change during 1*h* - *H*_2_*O*_2_ stimulation (Figure SI 2A). Assuming an average cytosolic pH of 7.1 for MCF7 in normal conditions [56], we estimated that pH decreased to an average of 6.9 at 55*min* after 100*μM H*_2_*O*_2_ exposure (Figure SI 2D). Importantly, even if pH variation exhibited cell-to-cell variability, pH did not drop down below pH 6.5 (Figure SI 2D). Second, we calibrated *R* variation for oxidized Grx1-roGFP2 in cells upon pH changes (see Supporting Information for details). We found no significant change in *R* by switching pH from 7.1 to 6.9 (Figure 2A-B). Furthermore, while *R* decreased upon prolonged exposure to *H*_2_*O*_2_, *R* tended to increase when exposed to pH 6.7 and 6.5 in our experimental conditions (Figure 2A-B). Finally, we compared variation of each fluorescent channels (under 420*nm* and 482*nm* excitation) upon exposure to *H*_2_*O*_2_ (Figure 2F-H) and pH change (Figure 2C-E). As expected, we found that fluorescence decreased in both channels when pH is decreased. On the other hand, signal in both channels increased upon *H*_2_*O*_2_ exposure. This demonstrates that the observed change in *R* under *H*_2_*O*_2_ stimulation cannot be attributed to pH variation in roGFP2 spectral properties.

**Figure 2:**
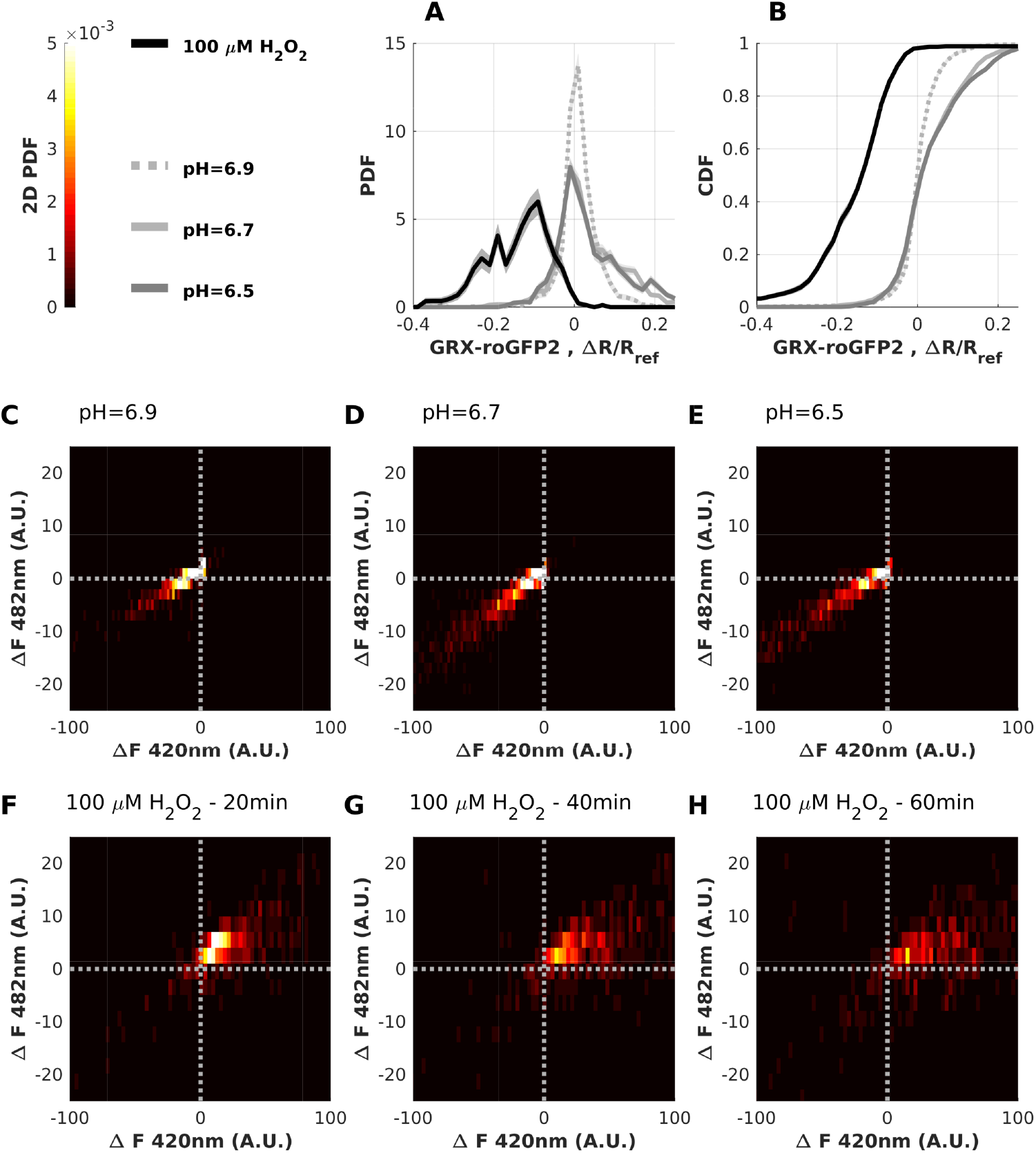
pH calibration of Grx1-roGFP2 confirms the existence of adaptative phenotype: MCF7 cells expressing cytoplasmic Grx1-roGFP2 were exposed to 100*μM H*_2_*O*_2_ in DPBS with 4.5g/L glucose for 10*min*. Then cells were fixed using 4% para-formaldehyde dissolved in DPBS (Sigma-Aldrich, L’Isle d’Abeau Chesnes, France). We finally exposed cells to various pH (7.1, 6.9, 6.7 and 6.5) and recorded corresponding variation of Grx1-roGFP2 fluorescence ratio. **A-B** - Statistics of variation of Grx1-roGFP2 ratio in its oxidized state, Δ*R*, relative to fluorescence ratio, *R_ref_*, in basal conditions (pH=7.1): **A** - Probability Density Function, PDF and **B** - Cumulative Density Function, CDF. For CDF the y-axis can be read as the probability to find a cell with a shape parameter higher than x-axis. For both PDF and CDF estimated from empirical distribution the gray shaded area represents 65% confidence interval obtained by bootstrap resampling [57]. **C-E** - Variation of each fluorescence channel upon pH change. For each cell, fluorescence variation, Δ*F*, is defined as the difference between fluorescence intensity at a given pH and for the reference pH. Cell distribution (bidimensional PDF) as a function of variation in both fluorescence channels, Δ*F*482*nm*, and Δ*F*420*nm* for: **C** - pH=6.9; **D** - pH=6.7; **E** - pH=6.5. **F-H** - Variation of each fluorescence channel at various time points following 100*μM H*_2_*O*_2_ addition (same data as Figure 1). Cell distribution (bidimensional PDF) as a function of variation in both fluorescence channels, Δ*F*482*nm*, and Δ*F*420*nm* for: **F** - *t* = 20*min*; **G** - *t* = 40*min*; **H** - *t* = 20*min*.

Variation of cytosolic pH could affect the Grx1-roGFP2 probe *via* another way. Indeed, both glutathione redox potential and the roGFP2 chromophore depend on the pH. This can in turn affect the probe’s degree of oxidation [53]. However, this effect is expected to be negligible because variation of the roGFP2 redox potential with pH compensates the one of glutathione [53]. Moreover, calibrating the Grx1-roGFP2 fluorescence ratio and using the pH estimation under *H*_2_*O*_2_ exposure, we could rigorously exclude this hypothesis (see supporting information for details). Indeed, we estimated that variation of cytosolic pH from 7.1 to 6.5 (a lower bound of pH in our experimental conditions) affects Grx1-roGFP2 fluorescence ratio by only 10^−4^% while we observed a drop by up to 40% on average upon prolonged *H*_2_*O*_2_ exposure.

Altogether, these results rule out a hypothetical side effect and highlight intracellular regulation that leads to a decrease of glutathione redox potential. This change in GSH redox potential reflects a gain in the cells reducing capacity. This regulation provides pseudo-adaptation phenotype for a significant number of cells (Figure 1E-F). Indeed for these cells, Grx1-roGFP2 signal increased shortly after stimulus addition but later decreased down even if extracellular *H*_2_*O*_2_ was kept constant. Interestingly signs of adaptation were also observed upon stimulus removal. Indeed for a significant number of cells we observed, upon removal, a drop of *R* signal below pre-stimulus level (Figure 1E). This drop was preserved for an hour post-stimulus removal. Evidence for adaptation upon stimulus addition was retained at the population averaged level (Figure 1I).

### Intracellular regulation is dose-dependent

To describe dynamic adaptation to *H*_2_*O*_2_ stimulus, we fitted Grx1-roGFP2 dynamics upon stimulus addition and removal. We defined 10 fitting parameters characterizing the temporal evolution *R*(*t*) (see Supporting Information). While some parameters pairs displayed strong linear correlation (see Figure SI 4) we were not able to reduce dimensionality using principal component analysis [58]. Indeed 7 parameters were necessary to explain 95% of the observed variance. However, we found strong linear correlation (with correlation coefficient, *r* > 0.81) between the first two fitting parameters, reflecting respectively the maximum signal variation, Δ*R_max_* = *R_max_* – *R*(*t* = 0), and the amount by which the signal was reduced after prolonged stimulation (see Figure SI 3). This suggests that the more the scavenging system is perturbed the stronger is the adaptation. Moreover we found high correlation (|*r*| > 0.7) between Δ*R_max_* and the signal variation upon stimulus removal Δ*R_rmv_*. This may indicate another sign of adaptation upon removal, which reflects the strength of intracellular change upon exposure to oxidative perturbation. Importantly, we did not find strong correlation between Grx1-roGFP2 expression level and parameters reflecting regulation dynamics (|*r*| < 0.27, see Figure SI 4). This suggests that the probe did not significantly perturb the effects described in this work. Moreover, apparent cell surface has low linear correlation coefficient (|*r*| < 0.18) with any of the parameters (Figure SI 3). While cell area in contact with extracellular medium will affect *H*_2_*O*_2_ import rate, it does not have a major influence on the dynamics reported here.

To gain understanding on the intracellular regulation described above, we repeated 1*h* step stimulation at various *H*_2_*O*_2_ concentrations ranging from 10 to 500*μM*. The magnitude of the stimulus influenced both the amplitude and the adaptation phenotype of the response (Figure 3A-B). Although there was a significant dependency of Grx1-roGFP2 dynamics with *H*_2_*O*_2_ concentration, our data also highlighted wide cell-to-cell variability. To describe single-cell dynamics quantitatively, we defined three empirical shape parameters which were schematically defined in Figure 3C-E. In order to account for both cell-to-cell variability and dose dependency of the shape parameters, we reported: (*i*) distribution of the parameters across the cell population (Figure SI 4A-D) and (*ii*) dependency of the most probable parameter value with extracellular *H*_2_*O*_2_ concentration, [*H*_2_*O*_2_] (Figure 3F-H). First, we noted that pre-stimulus fluorescence ratio, *R*_0_, was varying from cell-to-cell but its distribution was reproducible from one experiment to another (Figure SI 4A). Second, we examined the maximum variation of Grx1-roGFP2 fluorescence ratio, Δ*R_max_*, upon *H*_2_*O*_2_ addition (Figure SI 4B and Figure 3F). Δ*R_max_* distribution was clearly shifted toward higher values when [*H*_2_*O*_2_] was increased. This reflects a higher *H*_2_*O*_2_ effective production rate, Γ_*tot*_. Δ*R_max_* showed saturation for high [*H*_2_*O*_2_] and was well fitted by a Hill function with Hill coefficient of *n_H_* = 1 and half maximum concentration of [*H*_2_*O*_2_]_*H*_ = 70*μM* (Figure 3F). Such saturation might be due to saturation of the Grx1-roGFP2 probe itself [40].

**Figure 3:**
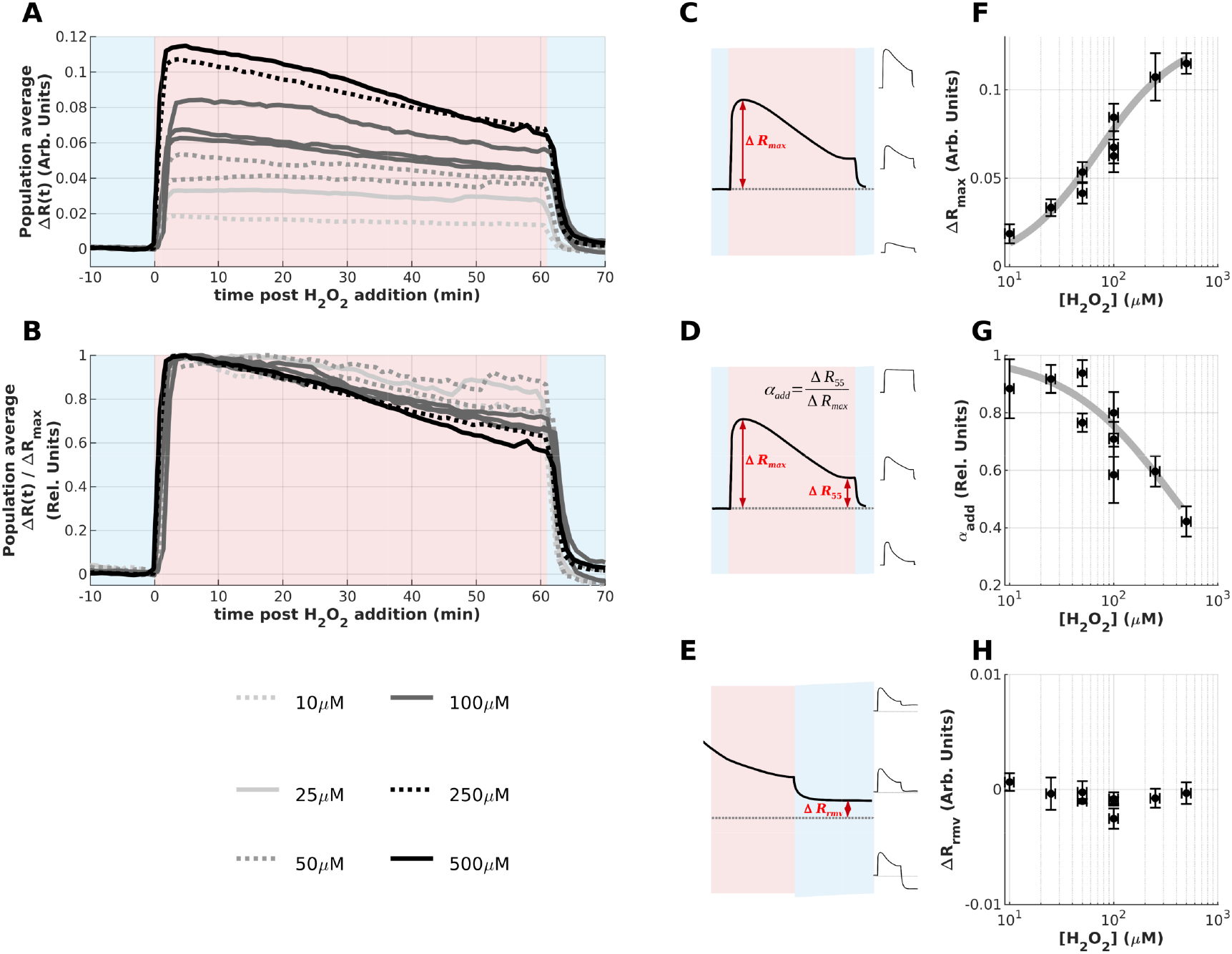
Influence of external *H*_2_*O*_2_ concentration on adaptation dynamics: We repeated experiments reported in Figure 1 at several *H*_2_*O*_2_ concentrations ranging from 10 *μM* to 500*μM*. To highlight both magnitude of the response and adaptative phenotype, we reported time evolution of both: **A** - absolute variation of *R*, Δ*R*(*t*) and, **B** - variation of *R* relative to its maximum Δ*R_max_*. Data are plotted with gray level representing *H*_2_*O*_2_ concentration: lighter gray corresponds to lower concentration and darkness is increasing with [*H*_2_*O*_2_] (see legend). In order to quantitatively describe dynamics of *R*(*t*) at the single-cell level we defined shape parameters: **C** - maximum Δ*R* upon stimulus addition (Δ*R_max_*); **D** - adaptation index (*α_add_*) defined as the ratio of Δ*R* 55*min* after addition and Δ*R_add_*; **E** - maximum Δ*R* upon stimulus removal (Δ*R_rmv_*). **F-G** - Dose-dependency of shape parameters - We reported dependency of the most probable shape parameter as a function of *H*_2_*O*_2_ concentration ([*H*_2_*O*_2_]) for **F** - Δ*R_max_*; **G** - *α_add_*; **H** - Δ*R_rmv_*. Vertical error bars represent 65% confidence interval obtained by bootstrap resampling [57] and horizontal error bars represent dilution error of 10% in our experimental conditions.

Then, we defined an adaptation index, *α_add_*, as the ratio between Δ*R*_55_ = *R*(*t* = 55*min*) – *R*(*t* = 0) and Δ*R_max_*. *α_add_* close to 0 characterize a near-perfect adaptation when the value of *R* at 55*min* post-stimulus was restored down to pre-stimulus level. On the opposite *α_add_* of 1 translated non-adaptive behavior when *R* stayed at its maximum 55*min* post-stimulation. Interestingly, *α_add_* decreased with [*H*_2_*O*_2_]: adaptative phenotype was favored for higher concentration (Figure SI 4C and Figure 3G). *α_add_* exhibited non-linear dependency with [*H*_2_*O*_2_] and was well fitted by a Hill function with Hill coefficient of *n_H_* = 0.8 and half-maximum concentration of [*H*_2_*O*_2_]_*H*_ = 390*μM* (Figure 3G).

Finally, we examined Δ*R_rmv_*, the difference between Grx1-roGFP2 fluorescence ratio after stimulus removal and pre-stimulus level *R*_0_. Δ*R_rmv_* showed large cell-to-cell variability (Figure 4D) but no evident dependency with [*H*_2_*O*_2_] (Figure 3H). Δ*R_rmv_* was negative for the majority of cells which reflected a lower Grx1-roGFP2 signal after removal compared to its pre-stimulus value. This suggests that glutathione reduced form GSH is favored after prolonged exposure to *H*_2_*O*_2_ compared to pre-stimulus. Such a difference corroborates the existence of adaptation on the ROS scavenging system.

**Figure 4:**
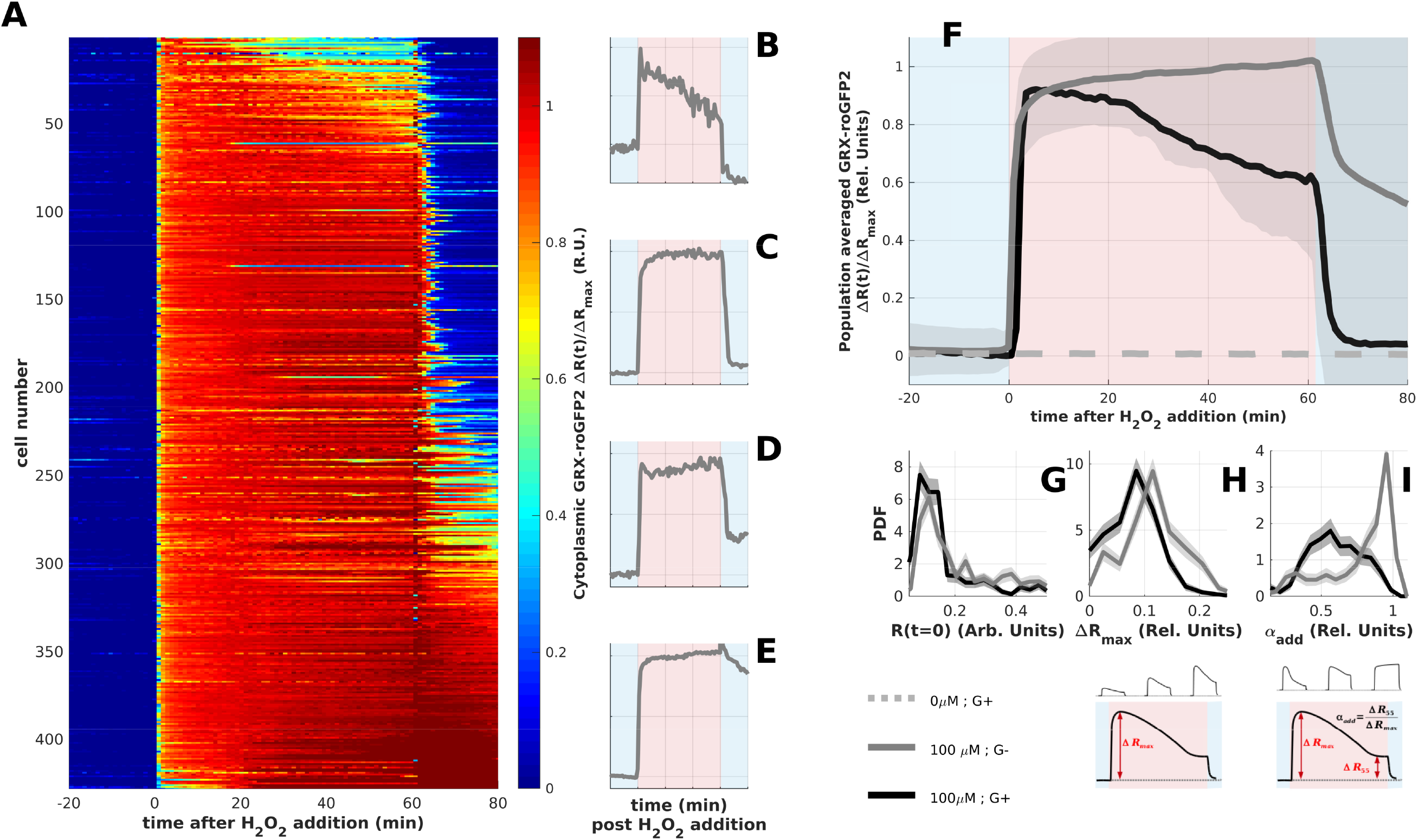
Adaptation depends on external glucose supply: Cells were exposed to *H*_2_*O*_2_ free DPBS with (or without) 4.5g/L glucose for 30*min* then exposed to 100*μM H*_2_*O*_2_ in DPBS with (or without) 4.5g/L glucose for 60*min* and recovered in *H*_2_*O*_2_ free DPBS with (or without) 4.5g/L glucose for another 60*min*. We imaged 435 cells at 1 frame per *min*. We plotted change in Grx1-roGFP2 fluorescence ratio compared to its pre-stimulus level, Δ*R*, relative to maximum change upon *H*_2_*O*_2_ addition Δ*R_max_*. **A** - false color image displaying, Δ*R*(*t*) / Δ*R_add_* as a function of time for all cells. **B** - Change in Grx1-roGFP2 signal averaged over the whole cell population for cells stimulated in the presence of extracellular glucose (black line) and cells stimulated without extracellular glucose (gray line) and for a control experiment in which *H*_2_*O*_2_ was replaced by mq-water (gray dashed line). Error bar represents standard deviation. **B-E** - Time traces of G+ single-cells: #1 (*α_add_* = 0.31), #113 (*α_add_* = 0.94), #225 (*α_add_* = 1.02), #337 (*α_add_* = 1.1). In order to quantitatively describe dynamics of *R*(*t*) we report shape parameters: **C** - initial state, *R*_0_; **D** - maximum Δ*R* upon stimulus addition (Δ*R_max_*); **E** - adaptation index (*α_add_*) defined as the ratio of Δ*R* 55*min* after addition and Δ*R_add_*. Lower panels show schematic definition of shape parameters. Are reported the Probability density Function (PDF, middle panel) for both conditions: with (black line) and without glucose (gray line). For both PDF from empirical distribution the gray shaded area represents 65% confidence interval obtained by bootstrap resampling [57].

### Glucose uptake limits adaptation via a NADPH-dependent mechanism

Glutathione and thioredoxin systems are two of the main intracellular ways for *H*_2_*O*_2_ (and reactive oxygen species) detoxification [59]. Both systems are well known to be powered by Nicotinamide adenine dinucleotide phosphate, NADP. In particular, its reduced form (NADPH) is necessary to reduce back GSSG and oxidized thioredoxins [18]. Moreover, NADPH is known to prevent oxidative inhibition of catalase [60, 61] a major enzyme catalyzing *H*_2_*O*_2_ degradation efficiently [62]. NADPH is produced by several glucose-dependent metabolic routes, including the pentose phosphate pathway (PPP) [18]. We thus examined the influence of external glucose on the adaptation dynamics.

We applied a 1*h* - 100*μM H*_2_*O*_2_ step stimulus to MCF7 cells expressing cytosolic Grx1-roGFP2 in the presence (G+) or absence (G-) of external glucose. Qualitatively, the Grx1-roGFP2 signal clearly showed that regulation was inhibited for G-cells (Figure 4A-E) compared to G+ ones (Figure 1D-H). On average, Grx1-roGFP2 ratio, *R*, decreased by ~40% in G+ cells (Figure 1D) while it kept increasing for G-cells (Figure 4F). While pre-stimulus Grx1-roGFP2 fluorescence ratios (*R*_0_, reflecting basal GSH/GSSG redox potential) were similar for both G+ and G-cases (see Figure 4G), other shape parameters were significantly affected by the absence of extracellular glucose. The maximum ratio variation (Δ*R_max_*) was increased by 20% on average in the absence of extracellular glucose (Figure 4H). Importantly the adaptation index, *α_add_*, was shifted from 0.6 on average in G+ cells to nearly 1 in G-cells (Figure 4I). Experiments with G-cells were repeated twice with similar results. These data suggest that external glucose is necessary to power the reported adaptation.

We then focused on the dynamics of NADPH concentration during and after *H*_2_*O*_2_ stimulation. For this purpose, we took advantage of NADP autofluorescence properties: its reduced form, NADPH, is fluorescent under UV excitation while its oxidized form, NADP^+^, is not [41]. We note that Nicotinamide adenine dinucleotide, NADH, also exhibit auto-fluorescence similar to NADPH. However NADH should not be affected by *H*_2_*O*_2_ load [18]. Moreover NADP is primarily located in the cytosol while NAD is located in the mitochondria[63]. Then, because fluorescence intensity was very different in the cytosol compared to mitochondria, the two subcellular compartments were easily separated by means of image processing.

Initial fluorescence level, *F*_0_, was clearly shifted toward lower values in the absence of extracellular glucose (Figure 5H). This suggests that NADPH level is lowered in G-cells. We applied a 1*h* - 100*μM H*_2_*O*_2_ step stimulus to wild type MCF7 cells while monitoring 460*nm* fluorescence under 365*nm* excitation in the cytosol. In the presence of extracellular glucose, cytosolic NADPH concentration showed very small changes upon *H*_2_*O*_2_ stimulation (see Figure 5A-D, 5G). Indeed one observed a small, but significant, drop (4% on average) shortly after stimulation followed by a fast re-increase (within 10min) even if the stimulus was maintained (Figure 5C-D, 5G). The observed undershoot was followed by a small and slow drift of NADPH fluorescence (Figure 5C-D, 5G). This drift is presumably due to pH variation which affects NADPH spectral properties [64]. In the absence of extracellular glucose the undershoot observed in G+ cells was even more pronounced in G-cells (Figure 5B, 5E-F, 5G). Indeed Δ*F_add_* relative to *F*_0_ was 50% in G-cells compared to 4% in G+ cells. However, we found that NADPH level stayed lower for G-cells compared to G+ under prolonged exposure to *H*_2_*O*_2_ (Figure 5B, 5E-F and 5I). Finally NADPH level was restored close to pre-stimulus level in both G+ and G-cells (Figure 5G). Experiments with G- and G+ cells were repeated twice with similar results. Altogether these results indicate that absence of extracellular glucose limits NADPH production which, in turn, limits regulation and adaptation.

**Figure 5:**
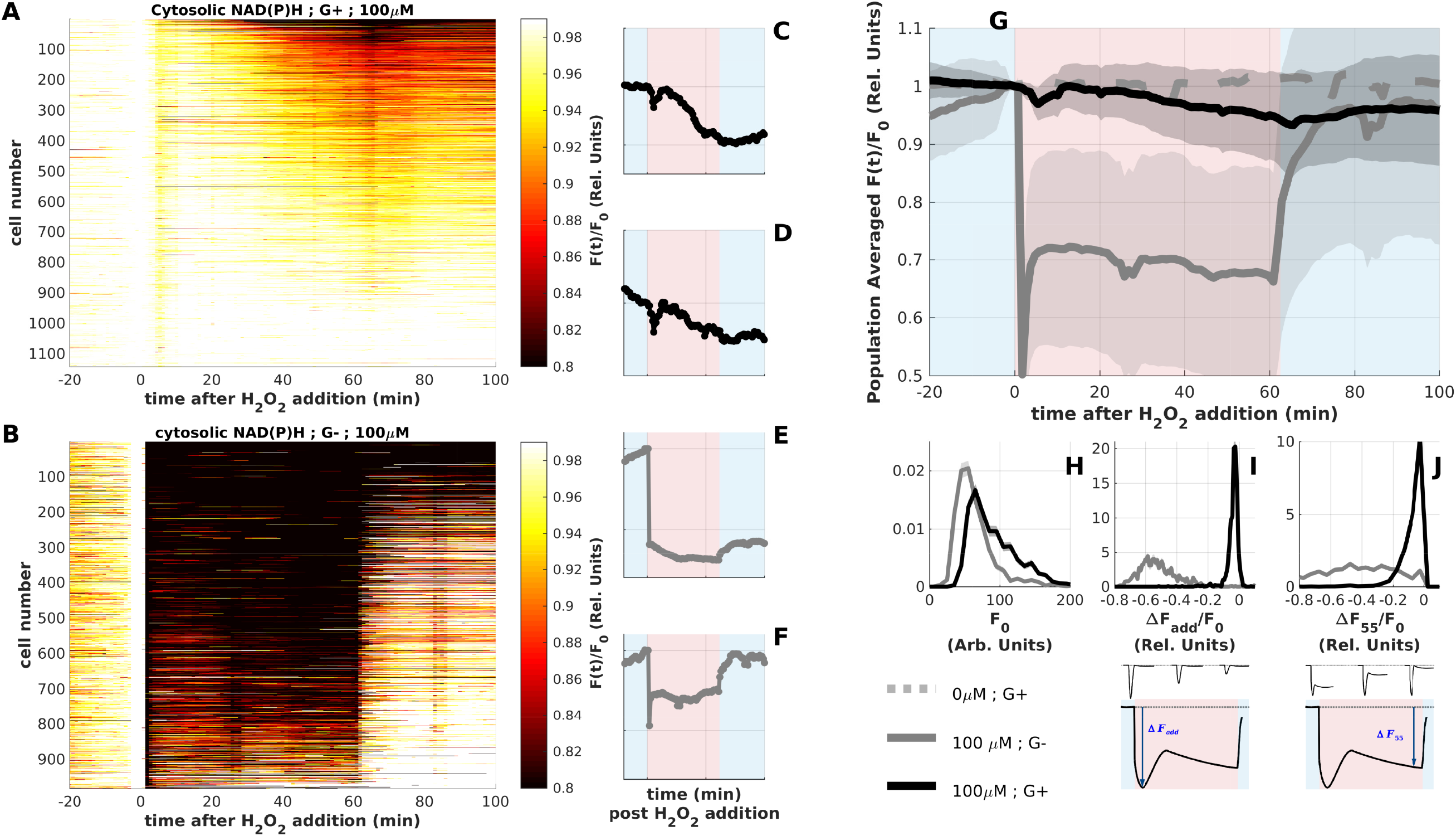
NAD(P)H dynamics associated with glucose-dependent adaptation: Cells were exposed to *H*_2_*O*_2_ free DPBS with (or without) 4.5g/L glucose for 30*min* then exposed to 100*μM H*_2_*O*_2_ in DPBS with (or without) 4.5g/L glucose for 60*min* and recovered in *H*_2_*O*_2_ free DPBS with (or without) 4.5g/L glucose for another 60*min*. We recorded cytoplasmic NAD(P)H autofluorescence (ex.365nm-em.460nm) in the presence (G+, black line) and absence (G-, gray line) of glucose. **A** - False color image displaying, NAD(P)H Fluorescence relative to its pre-stimulus level, *F*(*t*)/*F*_0_ as a function of time for all G+ cells. We imaged 1118 cells at 1 frame per *min*. **B** - False color image displaying, NAD(P)H Fluorescence relative to its pre-stimulus level, *F*(*t*)/*F*_0_ as a function of time for all G-cells. We imaged 987 cells at 1 frame per *min*. **C-D** - Time traces of G+ single-cells: #1 (Δ*F_add_*/*F*_0_ = 0.04 and Δ*F*_55_/*F*_0_ = 0.91), #561 (Δ*F_add_*/*F*_0_ = 0.04 and Δ*F*_55_/*F*_0_ = 0.96). **E-F** - Time traces of G-single-cells: #1 (Δ*F_add_*/*F*_0_ = 0.41 and Δ*F*_55_/*F*_0_ = 0.30), #481 (Δ*F_add_*/*F*_0_ = 0.53 and Δ*F*_55_/*F*_0_ = 0.75). **G** - Population average of *F*(*t*)/*F*_0_ error bars represent standard deviation. G+ cells were represented in black, G-cells were reported in gray and control experiment in which *H*_2_*O*_2_ was replaced by mq-water was depicted in gray dashed line. Error bar represents standard deviation. **H-J** - Probability density function of shape parameters. Lower panels shows schematic definition of shape parameters. Grey shaded area represents 65% confidence interval obtained by bootstrap resampling [57]. G+ cells are in black and G-cells are in gray. **H** - probability density of pre-stimulus NAD(P)H autofluorescence level, *F*_0_, reflecting basal NAD(P)H concentration. **I** - probability density of maximum variation in NAD(P)H autofluorescence relative to *F*_0_ (Δ*F_add_*/*F*_0_). **J** - probability density function of variation in NAD(P)H autofluorescence 55 min post-stimulation relative to *F*_0_ (Δ*F*_55_/*F*_0_).

## DISCUSSION AND CONCLUSIONS

In this work, we have used live cell imaging together with controlled *H*_2_*O*_2_ stimulation to monitor dynamic adaptation to oxidative load. Our data suggested at least two types of regulations. One faster regulation was reflected by NAPDH dynamics. Upon stimulus addition, NAPDH-level rapidly dropped but was restored to pre-stimulus level within ten minutes. This behavior has been reported by others using time-resolved metabolomics experiments [15] or using a NADPH fluorescent reporter [25]. The second regulation was slower and, to our knowledge, has never been observed as such on the ROS scavenging system. This slower regulation led to a change of glutathione redox potential upon prolonged *H*_2_*O*_2_ stimulation: while reducing capacity of the glutathione system rapidly decreased upon *H*_2_*O*_2_ addition, we found that this reducing capacity was slowly restored even if the stimulus was maintained. As GSH serves for *H*_2_*O*_2_ scavenging but, more generally, for peroxides reduction including protein detoxification [16, 17] Grx1-roGFP2 can be seen as a readout of cellular reducing capacity.

One question immediately arises: what are the molecular networks underlying the observed adaptations? One hypothesis is that rerouting of metabolic flux leads to an increase of reduction rate of NADPH and GSH cofactors. Let us first consider this hypothesis in light of recent reports that have provided insights toward deciphering regulations and adaptative response of PPP during oxidative load. Glucose-6-Phosphate deshydrogenase (G6PDH) is one of the major NADPH producing enzyme and regulates the flux entering the PPP[16, 17]. G6PDH is known for a long time to be far from saturation in basal conditions [65]. This allows the so called reserve flux capacity of PPP that is provided by NADPH self inhibition of its own production *via* G6PDH [66]. This allosteric inhibition was shown to allow extremely rapid flux rerouting from glycolysis toward the PPP[14, 19]. However effects of this regulation occur in less than a minute after oxidative load and thus cannot explain the slower Grx1-rogFP2 nor the NADPH signals observed in this work. Later other regulations allow a further increase of flux into the PPP. Kuehne *et al*. have reported recycling of carbon from the oxidative branch of the PPP toward glycolysis. Such a rerouting in turn reverses the glycolytic flux and leads to a further increase of G6PDH activity within ten minutes[15]. This time scale is compatible with the dynamics we observed for NADPH. This observation supports an hypothesis in which NADPH level drops just after stimulus addition but is later restored *via* increased G6PDH activity. However the time scales involved for adaptation of GSH redox potential are much slower than these metabolic effects.

Another important observation of this work is that the presence of external glucose was necessary for adaptation of GSH redox potential. Moreover, the absence of external glucose prevented from restoration of NADPH to pre-stimulus level. This suggests that glucose uptake limits adaptation of GSH redox potential by limiting the flux through the oxidative PPP. This was expected as GSH reduction is powered by NADPH, which is itself produced *via* the PPP. However, we note that it does not mean that the regulation responsible for the slow adaptation of GSH redox potential is glucose-dependent. External glucose can be necessary for adaptation but not sufficient. Slower processes such as transcriptionnal activity [9, 10] and GSH *de novo* synthesis [13] have to be considered to explain the adaptation of GSH redox potential. Indeed Grx1-rogFP2 is sensitive to total glutathione concentration as increased GSH level would affect its redox potential. Such an increase of total GSH would enhance the cell reducing capacity.

The adaptation index, *α_add_*, allowed us to assess adaptation efficiency at the single-cell level. Indeed *α_add_* reflects the residual modification of cellular redox state (Δ*R_rmv_*) relative to the magnitude of the perturbation (Δ*R_max_*). But it is also a readout of the adaptation rate: *K* ∝ 1 – *α_add_*. Increasing *H*_2_*O*_2_ concentration clearly induced a transfer from non-adapting phenotypes to adapting ones (see Figure SI 4C). Measurement of dose-response revealed an activation threshold for triggering adaptation of GSH redox potential. Such threshold was described in this work by fitting a sigmoidal function (see Figure 3G, [*H*_2_*O*_2_]_*th*_ ~ 70*μM*). Thresholds are known features of several network motifs [67] and are consistent with activation of redox-sensitive factors [10].

It can appear surprising that the Grx1-roGFP2 signal did not display an echo of the NADPH under-shoot upon stimulus addition. Let us here stress several aspects that could explain this observation. First glutathione system is at the entry of the ROS scavenging system. It is indeed directly coupled to *H*_2_*O*_2_ *via* Glutathione Peroxidase. On the other side, NADPH is coupled to both glutathione *via* glutathione reductase and to the large metabolic network. It is then plausible that GSH dynamic exhibits less inertia than NADPH dynamics, and an overshoot on Grx1-roGFP2 signal, if it exists, may be faster responding than the NADPH undershoot. Second the probe’s rise and decay times are both of the order of 1 min, and the rise time of the stimulus in our experimental conditions is also of one minute. Then our time resolution may not have been sufficient to observe a fast Grx1-roGFP2 time evolution. One observation may corroborate this hypothesis. In some experiments, we have observed a small overshoot upon stimulus addition that involved at most one time-point. This effect was varying from cell-to-cell and was not reproducible from one experiment to another. Improvements in our fluidic system and increasing the recording rate may help to better resolve such an overshoot on Grx1-roGFP2 signal (if it exists) in a near future.

Even if single information has not been thoroughly exploited in this work, our data yet revealed high cell-to-cell variability in response. Such heterogeneity in a clonal cell line can arise from variation in proteome [51, 68, 69]. It was shown using mathematical modeling of both stress response network and death decision pathway that cell-to-cell variation and noise in signaling networks can lead to fractional killing [70, 71, 72]. It is tempting to ask whether and how the cell-to-cell variability that we observed here affects cell death or survival. Are the cells that exhibit better adaptation to the stimulus the same as the cells that survive? Answer to this a question may not be as trivial as one may expect. Recent reports have indeed unveiled that intracellular ROS level does not correlate with cell fate [73]. Although our experimental framework allows to record time evolution of less species compared to time-resolved metabolomics experiments, our system enables live cell imaging together with high-throughput single-cell data. It will be well suited to follow simultaneously dynamics of the ROS scavenging system and cell death modules. Moreover we note that our methodology is not limited to GSH redox potential nor NADPH level. Our protocol can be used to monitor a variety of recently developed fluorescent redox probes [55, 40, 50, 47, 49, 74, 25, 53, 75]. These probes are now engineered in many different colors [75, 49, 53]. This will allow one to correlate single-cell dynamics of several species involved ROS scavenging with fluorescent reporters of cell death decision pathways [76].

## MATERIALS AND METHODS

### Cell culture

MCF7 cancer cell line was purchased from the American Type Culture Collection (ATCC, Manassas, VA). These adherent cells are grown as monolayer in Dulbecco’s modified Eagle’s medium (DMEM; Lonza, Levallois-Perret, France) supplemented with 10% (v/v) fetal bovine serum (FBS; Life Technologies, Saint-Aubin, France), 1% L-glutamine (2 mM) and 1% (v/v) penicillin-streptomycin (100 IU/ml) (Lonza). Cell cultures are maintained at 37°C in a humidified atmosphere containing 5 % CO_2_ (v/v), and passage at preconfluence (twice a week) using 0.05 % trypsin-0.53 mM ethylenediamine tetraacetate (EDTA; Lonza). MCF7 growing cells are routinely screened for the presence of mycoplasma using DNA-staining with the nuclear dye Hoechst 33342 (1:10000 dilution) (Sigma-Aldrich, L’Isle d’Abeau Chesnes, France) to avoid collecting data from unknowingly contaminated cell cultures.

### Cell transfection

Experiments monitoring cytosolic Grx1-roGFP2 [40] probes were performed on stably transfected cell lines. For stable cell line transfection, Wild-type MCF7 cells (MCF7 WT) were transfected with a plasmid expressing the sensor of interest using FuGENE HD transfection reagent (Promega, Charbonnières, France) according to the manufacturer’s instructions. The stable transfected cell line was then established under selective pressure by 1000 μg/ml geneticin (Life Technologies). pEIGW Grx1-roGFP2 was a gift from Tobias Dick (Addgene plasmid # 64990; http://n2t.net/addgene:64990; RRID:Addgene_64990). The Grx1-roGFP2 sequence was cloned into a pEGFP-N1 backbone (Clontech) with CMV promotor to avoid lentiviral transfection strategy. For this we use the In-Fusion PCR cloning system (Clontech) according to the manufacturer’s instructions. Grx1-roGFP2 sequence was clived at Xma1 and Not1 restriction sites and EGFP was clived at BamH1 and Not1 restriction sites.

### Fluidic chamber preparation

Fluidic chambers were built by assembling a 25mm × 75mm glass coverslip #1.5 (#10812, Clinisciences, Nanterre, France) and a 1.6mm thick silicone sheet (Red Silicone Sheet, Grace Bio-Labs). The silicone sheet was cut to fit the 25mm × 75mm glass coverslip, and a 5mm × 50mm channel was punched with a custom-made tool. Glass and silicone do not need adhesive to build the chamber. They stick together *via* electrostatic interactions. Cells were seeded on a first half of the chamber (without the top slide) 48h prior to imaging. Cells were cultured as traditional culture dishes at 37°C in a humidified atmosphere containing 5 % CO_2_ (v/v). For this, we placed several chambers in 150mm diameter culture dish (#83.3903, Sarstedt, Marnay, France). Chambers were raised using custom made holders and the 150mm diameter culture dish was filled with 30mL sterile water to avoid medium evaporation in the chamber. All experiments were performed on at least 2-days-old cell cultures (50 % final confluence) prepared in complete DMEM. The chamber was sealed just before imaging. For this a polymer slide with 0.8mm height channel and fluidic connections (Ibidi, sticky-Slide I Luer #80168) was placed on top. Again, slide and silicone do not need adhesive to build the chamber. They stick together *via* electrostatic interactions.

### Live cells imaging

For imaging we maintained cells in Dulbecco’s Phosphate Buffered Saline with *MgCl_2_* and *CaCl_2_* (DPBS; #D8662-500ML, Sigma-Aldrich, L’Isle d’Abeau Chesnes, France). DPBS was supplemented with 4.5g/L glucose (D-(+)-Glucose, #G7021, Sigma-Aldrich, L’Isle d’Abeau Chesnes, France) except when otherwise stated. DPBS solutions were prepared in 50mL tubes and placed in dry heating bath at 37°C under 5 % CO_2_ (v/v) humidified atmosphere. DPBS containing *H*_2_*O*_2_ was prepared 1h prior to the experiment from a 10*mM* solution. Concentrated (10*mM*) solution was prepared on ice the day of the experiment by adding 10*μL* of stock *H*_2_*O*_2_ solution 30% w/v (#16911, Sigma-Aldrich, L’Isle d’Abeau Chesnes, France) in 10mL sterile mq-water. Concentration of stock solution was verified daily by measuring 265nm Optical Density using a custom-made system.

After chambers were sealed (see above), samples are placed on a Nikon TiE microscope within 5-10min prior to experiment. Custom built imaging system was described previously[51]. The microscope with motorized filter wheel is equipped with a XY-motorized stage (ASI). Cells were imaged through a 60X microscope objective (NA=1.4, Nikon) on a sCMOS camera (OrcaFlash LT, Hamamatsu). We set the camera binning to 2 resulting in an effective pixel size of 325 nm. Illumination for fluorescence and brightfield imaging is achieved through custom built optical system (components from Thorlabs). We use LED light source (Thorlabs) for synchronization of illumination with other apparatus. Exposure time is set to 150ms for all experiments and for each fluorescence channel (as well as brightfield illumination). Light power density, filters set and LED for each type of experiments are summarized in supporting information (SI Table 1). We use a custom-built acquisition software written in Labview to control the setup. In order to increase the output rate of the experiment we acquire data for 40 different fields of view in the same sample (by use of the motorized stage) leading to the tracking of approximately 500 cells per experiment. Two consecutive fields of view are separated by approximately 100*μm*. Focusing is maintained *via* Nikon Perfect Focus System.

### Image processing and analysis

All image processing and data analysis were performed using custom written algorithms in Matlab. Briefly, we first acquired fluorescence images of dishes filled with fluorescent dyes (rhodamine 110 for SypHer and Grx1-roGFP2 and courmarin for NAD(P)H autofluorescence) for flat field correction. As cells do not move significantly during the time course of the experiment, we used a single image (time averaged fluorescence image: ex.482nm/em.520nm) for cell segmentation. Centers of cells were first selected manually. This allows us to separate neighboring cells by the perpendicular bisector of the cells centers. Then images were segmented using a modified Otsu thresholding method [77].

Estimation of Grx1-roGFP2 [40] fluorescence ratios was performed as follows: Flat field correction was applied to each image of the time series for both channel 2 (ex.482nm/em.520nm) and channel 1 (ex.430nm/em.520nm). A constant background was subtracted to each channel before further analysis. We estimated background by calculating median gray level of the region without cells in the image. A fluorescence ratio image was then computed by dividing each pixel of channel 2 by corresponding pixel of channel 1. Fluorescence ratio for each cell, *R*, was then obtained by averaging the ratiometric image over the corresponding cell mask. *R* exhibit slow decay over the time course of the experiment. This decay is due to slow interaction of the fluorophore with light [50] populating the probe’s dark states. We correct for baseline drift by fitting data with bi-exponential function for each single-cells time-series. We only used the 20 first frames for which cells were not exposed to stimulus. The fitted theoretical function was then substracted to raw data.

Estimation of NAD(P)H autofluorescence [41, 42] was performed as follows: Flat field correction was applied to each image of the time series of channel 0 (ex.365nm/em.460nm). A constant background was subtracted to each channel before further analysis. Constant background was estimated by fitting a truncated histogram of all images with a normal distribution. The histogram was truncated to limit high intensity pixels corresponding to NAD(P)H fluorescence. Cell segmentation was performed first as described above. Then cytosolic and mitochondrial signals were separated by applying Otsu thresholding method [77] a second time separately for each cells. This way we can split the cell mask in two regions corresponding to mitochondria separated from cytosol and nucleus. Cytosolic auto-fluorescence, *F*, was then computed for single-cells by averaging images over the corresponding cell mask. Each single-cell time series were normalized to initial fluorescence level *F*_0_. *F* exhibits a slow drift due to photo-bleaching of the sample. Baseline drift was corrected by fitting a single exponential decay to signal obtained for non-stimulated cells. Fitted theoretical function was then substracted to raw data.

## Supporting information

SIvideo1

## Acknowledgements and fundings

This work has been partially supported by the LABEX CEMPI (ANR-11-LABX-0007), as well as by the Ministry of Higher Education and Research, Hauts de France council and European Regional Development Fund (ERDF) through the Contrat de Projets Etat-Region (CPER Photonics for Society P4S).

We thank the PhLAM laboratory technical support and in particular: Herve Damart for electronics to control the setup, Gauthier Dekyndt for mechanical devices, Raoul Torero for daily support on the biology platform and Jean Pesez for daily assistance on instrumentation.

## Author Contributions

Performed experiments: D.S. F.D. F.A.; Assisted with experiments: A.P. M.G. M.B. C.L.; Designed research B.P. Q.T. E.C. F.A.; Analyzed data: F.A.; Wrote the main text of the manuscript: F.A. All authors have read, commented and approved the final version of the manuscript.

## Declaration of competing interest

The authors declare that they have no known competing financial interests or personal relationships that could have appeared to influence the work reported in this paper.

## Notes

### Competing Interest Statement

The authors have declared no competing interest.

