## Supporting Information for "Measuring adaptation dynamics to hydrogen peroxide in single human cells using fluorescent reporters"

Dana Simiuc[1], Fatima Dahmani[1], Alexandra Pruvost[1], Marie Guilbert[1], Mathilde Brulé[2], Chann Lagadec[2], Quentin Thommen[1], Benjamin Pfeuty[1], Emmanuel Courtade[1] and François Anquez[1*]

November 14, 2020

1 Univ. Lille, CNRS, UMR 8523 – PhLAM – Physique des Lasers Atomes et Molécules, F-59000 Lille, France

2 Univ. Lille, CNRS, UMR9020, Inserm, UMR-1277 – CANTHER – CANcer Heterogeneity, Plasticity and Resistance to THERapies", F-59000 Lille, France

### Hydrogen peroxide stability in various culture media

In order to highlight a potential feedback we need to apply a steady stimulus. Extracellular $H_2O_2$ is known to be degraded by culture medium and, furthermore, in the presence of cells[]. We monitored kinetics of $H_2O_2$ degradation in various media (DMEM, DPBS) in the presence or absence of cells. For this we used the commercial chemical probe HyPerBlu (Lumigen, Michigan, USA). HyPerBlu is an extracellular substrate that reacts directly with $H_2O_2$ to produce a luminescent product. Luminescence intensity was used to measure relative concentration of extracellular $H_2O_2$. We have used a plate-reader (FluoStar Omega, BMG LabTech, Champigny s/Marne, France) to detect time-resolved luminescence. We note that the plate-reader allowed to maintain samples at 37°C in humified and 5% $CO_2$ atmosphere.

For media without cells we proceed as follows : Medium under study was pre-warmed in the plate-reader. Cells were cultured in 3.5cm culture dish. At the beginning of the experiment we added $H_2O_2$ from stock solution (10$mM$ in mq-water) to obtain a 100$\mu M$ final $H_2O_2$ concentration. At given time point 96-well plate (ThermoFisher Scientific) was filled with 50$\mu L$ of the medium in each well. We used the plate-reader injectors to add 50$\mu L$ of HyPerBlu solution at given time points. Rapid kinetics of luminescence was monitored during 2min. We used the maximum of the luminescence as a measure of $H_2O_2$ concentration. Indeed calibration experiments showed that this quantity is proportional to $H_2O_2$ concentration. The operations was repeated at several time points to monitor, at slower time scale, the time evolution of $H_2O_2$ concentration (Figure SI 1A-C). 10 replicates were done for each concentrations and each media. Using these data we found that $H_2O_2$ concentration decays to 75% of initial concentration in complete DMEM without Pyruvate and to 60% in DMEM with Pyruvate 6 hours after injection (Figure SI 1A). Importantly we did not found significant decrease of $H_2O_2$ concentration DPBS in the absence and nor in the presence of 4.5g/L glucose even after 17h (Figure 1C).

For media in the presence of cells we proceed as follows : Cells were cultured at various confluences in complete DMEM. At various time points we withdraw 50$\mu L$ of extracellular medium from the culture dishes. This volume was placed in a well and mixed with 50$\mu L$ of HyPerBlu. We monitored luminescence to estimate $H_2O_2$ concentration. We found that $H_2O_2$ decay faster in the presence of cells and that decay rate increases with cell confluence (Figure SI 1B). This due to consumption of $H_2O_2$ by cell scavenging system.

### Estimation of pH variation upon hydrogen peroxide exposure

Cytosolic pH was shown to drop during $H_2O_2$ exposure [1]. Most fluorophores are pH sensitive. This raised questions on whether the observed change of Grx1-roGFP2 signal have to be attributed to pH change or to modification of glutathione redox potential due to an intracellular regulation. To answer this question we first needed to estimate the amplitude of pH change upon prolonged $H_2O_2$ stimulation.

For this purpose we used SypHer, a fluorescent pH probe [2, 3]. SypHer has two forms. A deprotonated form which emits green light ($520nm$) under $482nm$ excitation and a protonated form which emits green light ($520nm$) under $420nm$ excitation [3]. Here the quantity of interest is the fluorescence ratio ($482nm$ versus $420nm$),$R$. We note that definition of ratio for SypHer was the inverse of the one used Grx1-roGFP2. In the following we denote $C_1$ the concentration of the protonated form (excitation $420nm$ and $C_2$ the concentration of the deprotonated form (excitation $482nm$). The fluorescence ratio then reads :

$$R = \frac{\sigma_2 \times \Phi_2 \times \phi_2 \times I_2 \times C_2}{\sigma_1 \times \Phi_1 \times \phi_1 \times I_1 \times C_1} \tag{SI 1}$$

where $\sigma_i$ is the absorption cross section, $\Phi_i$ the quantum efficiency, $\phi_i$ a coefficient reflecting apparatus transmission and $I_i$ the excitation intensity corresponding to form $i$.

Considering a reference pH with SypHer fluorescence ratio, $R_{ref}$, we defined the quantity :

$$\frac{\Delta R}{R_{ref}} = \frac{R - R_{ref}}{R_{ref}} = \frac{C_2}{C_1} \times \frac{C_1^{ref}}{C_2^{ref}} - 1 \tag{SI 2}$$

where $C_i^{ref}$ is concentration of form $i$ at the defined reference pH.

We note that $\frac{\Delta R}{R_{ref}}$ is independent of experimental conditions such as illumination intensity or apparatus. Indeed this quantity only depends on both reference pH and pH at which $\Delta R$ was measured. Thus calibrating $\frac{\Delta R}{R_{ref}}$ as a function of pH and taking cytosolic pH as a reference allowed us to estimate intracellular pH variation from SypHer measurements. Cytosolic pH is close to 7.1 in normal conditions [4]. In the following we used $pH = 7.1$ as reference.

We next performed $1h$ - $100\mu M$ $H_2O_2$ step stimulation experiments with cells expressing cytosolic SypHer. While seeding them in our fluidic chamber (48h prior to imaging), MCF7 WT cells were transiently transfected with plasmid encoding for SypHer with FuGENE HD transfection reagent (Promega, Charbonnières, France) according to the manufacturer's instructions. SypHer was a gift from Nicolas Demaurex (Addgene plasmid # 48250 ; `http://n2t.net/addgene:48250` ; RRID:Addgene_48250). Cells were then imaged while exposed to $1h$ - $100\mu M$ $H_2O_2$ step stimulus. Stimulation and imaging conditions were the exact same as for experiments reported in the main text with Grx1-roGFP2 probe. As expected, pH decayed following stimulation with $100\mu M$ $H_2O_2$ (Figure SI 2A). $\frac{\Delta R}{R_0}$ could be measured $55min$ post-stimulation (Figure SI 2B). Here $R_0$ denotes SypHer fluorescence ratio at the beginning of the experiments which we expect to correspond to $pH = 7.1$ (reference).

We next used data from [2] to calibrate SypHer $\frac{\Delta R}{R_{ref}}$ with reference pH of 7.1 (Figure SI 2C). Data were well fitted by Hill function :

$$\frac{\Delta R}{R_{ref}} = \frac{\Delta R_1}{R_{ref}} + \frac{\Delta R_2/R_{ref}}{1+e^{\nu(pKa-pH)}} - 1 \tag{SI 3}$$

where $R_{ref} = 0.758$ is the SypHer ratio at reference pH [2], $\Delta R_1 = 0.1$ and $\Delta R_1 = 9.5$ are constants, Hill coefficient was found to be $\nu = 2$ and we took $pKa = 8.4$.

Given this calibration, it was straightforward to use measured $\frac{\Delta R}{R_0}$ (Figure SI 2B) to estimate intracellular pH at $55min$ post $H_2O_2$ addition (Figure SI 2D). We found that, under $1h$ - $100\mu M$ $H_2O_2$ step stimulation, pH varied from 7.1 to 6.9 on average.

### Dependency of Grx1-roGFP2 spectral properties with pH change

We found that Grx1-roGFP2 ratio reached a maximum and decayed after prolonged $100\mu M$ $H_2O_2$ stimulation (Figure 1D-I and Figure 2A-B). Although Grx1-roGFP2 ratio, $R$, is pH insensitive close to physiological conditions [5], $R$ could be pH dependent in extreme cases because the spectral properties of the oxidized and reduced forms of Grx1-roGFP2 have different pH dependency[6]. We thus needed to test whether the amount of pH change estimated above could have led to significant variation of GRx1-roGFFP2 fluorescence ratio.

For this purpose, MCF7 cells stably expressing GRx1-roGFP2 were cultured in our fluidic chamber as explained in the main text. Cells were exposed to $100\mu M$ $H_2O_2$ in DPBS with glucose for $5min$. This time was sufficient to reach maximum signal of Grx1-roGFP2. Then cells were fixed using 4% para-formaldehyde dissolved in DPBS (Sigma-Aldrich, L'Isle d'Abeau Chesnes, France). We finally used our fluidic system to alternatively expose cells to various pH (7.1, 6.9, 6.7 and 6.5) and to record corresponding variation of GRx1-roGFP2 fluorescence ratio. We found no change in $R$ by switching from $pH = 7.1$ to $pH = 6.9$ (Figure 2A-B). Furthermore we found that $R$ slightly increases when pH is switched from 7.1 to 6.7 or to 6.5(Figure 2A-B) while $R$ decreased (after reaching a maximum) upon prolonged $H_2O_2$ exposure (Figure 1D-I and Figure 2A-B).

To further confirm that the observed changes of GRx1-roGF2 fluorescence ratio under $100\mu M$ $H_2O_2$ stimulation cannot be attributed to pH we examined variation of each fluorescence chanel separetely. As expected fluorescence intensity in both chanels decreased when pH decreases (Figure 2C-E). On the other hand $R$ increased in both chanel upon prolonged $H_2O_2$ (Figure 2F-H).

Altogether these results show that pH change is not responsible for the observed changes in $R$ during oxidative stress.

### Calibration of GRx1-roGFP2

pH could affect GRx1-roGFP2 not only *via* the spectral properties of the probe. Indeed redox potentials of both glutathione and the roGFP2 fluorophore depends on pH. Thus the efficiency of electron transfer from glutathione to roGFP2 depends on pH. As the dependency of both redox potential with pH are collinear (for pH below 8) the effect is expected to be small [6]. However, in order to rule out this effect rigorously we performed calibration experiments to estimate how such a pH effect would affect fluorescence ratio.

We first determined how the roGFP2 degree of oxidation, $D$, affects the Grx1-roGFP2 fluorescence ratio, $R$. roGFP2 degree of oxidation, $D$, is defined as the concentration of oxidized roGFP2 relative to roGFP2 total concentration :

$$D = \frac{[roGFP2_{ox}]}{[roGFP2_{ox}] + [roGFP2_{red}]}$$

Given the fluorescence ratio for the fully reduced probe, $R_{red}$, and the fluorescence ratio for the fully oxidized probe, $R_{ox}$, and an instrument factor, $\Phi$, the degree of oxidation reads [6] :

$$D = \frac{R - R_{red}}{\Phi\left(R_{ox} - R\right) + R - R_{red}}$$

where the instrument factor is defined as the ratio of fluorescence intensity in the $482nm$ excitation chanel of the fully oxidated probe, $I_{ox}$, and the fully reduced probe, $I_{red}$ :

$$\Phi = \frac{I_{ox}}{I_{red}}$$

We then derived dependency of $R$ with the oxidation degree :

$$R = \frac{D\left(\Phi R_{ox} - R_{red}\right) + R_{red}}{1 - D\left(1 - \Phi\right)} \tag{SI 4}$$

To estimate how a change in the oxidation degree will affect the fluorescence ratio, we calculated the derivative of expression SI 4 :

$$\frac{dR}{dD} = \Phi\frac{R_{ox} - R_{red}}{\left(1 - D(1 - \Phi)\right)^2}$$

The maximum influence of $D$ on the fluorescence ratio $R$ occurs for the highly oxidated probe :

$$\left.\frac{dR}{dD}\right|_{max} = \frac{R_{ox} - R_{red}}{\Phi} \tag{SI 5}$$

We next performed calibration experiments to estimate, $\Phi$, $R_{ox}$ and $R_{red}$. For this purpose, MCF7 cells stably expressing Grx1-roGFP2 were seeded in our fluidic chamber. While monitoring fluorescence in both chanels, we applied successively high oxidative load and highly reducing agent to estimate the parameters $\Phi$, $R_{ox}$ and $R_{red}$. We used $1mM$ $H_2O_2$ to monitor spectral properties of fully oxidized Grx1-roGFP2. We found that $500\mu M$ dithiothreitol (DTT) was sufficient to convert all GRx1-roGFP2 in its reduced form. $\Phi$

was estimated to be $0.49 \pm 0.06$ in our experimental conditions. Grx1-roGFP2 dynamic range, $R_{ox} - R_{red}$, varies from cell-to-cell in our experimental conditions. However we found the maximum dynamic range over the cell population to be smaller than 0.3 : $(R_{ox} - R_{red})_{max} < 0.3$

We now need to evaluate how a change in redox potential will affect the oxidation degree of GRx1-roGFP2. The relationship between roGFP2 oxidation degree, $D$ and glutathione redox potential, $E$, is described by the following equation [6] :

$$D = \frac{1}{1 + e^{-\frac{2F}{RT}(E - E_0)}}$$

where $E_0$ is the midpoint roGFP2 redox potential ($-280mV$), $F$ is the Faraday constant ($96.485C.mol^{-1}$), $R$ is the gas constant ($8.315J.K^{-1}.mol^{-1}$) and $T$ the absolute temperature ($310.15K$).

To estimate how a change in $E$ will affect oxidation degree, we calculated the derivative of this expression :

$$\frac{dD}{dE} = \frac{2F}{RT} \frac{e^{-\frac{2F}{RT}(E-E_0)}}{\left(1 + e^{-\frac{2F}{RT}(E-E_0)}\right)}$$

This expression is maximum for $E = E_0$ :

$$\left.\frac{dD}{dE}\right|_{max} = \frac{F}{2RT} \tag{SI 6}$$

Because variation with pH of roGFP2 and glutathione redox potentials deviate from collinearity, a decrease of cytosolic pH from 7.1 to 6.5 mimics reduction of glutathione and a decrease of glutathione redox potential, $E$. From data reported in **??**, we estimated that an upper bound for the amplitude of this effect on apparent glutathione redox potential is $\Delta E = 0.1mV$. Apparent variation of $E$ turns into a decrease of Grx1-roGFP2 fluorescence ratio. A upper bound for such a decrease reads :

$$\Delta R_{pH} = \left.\frac{dR}{dD}\right|_{max} \times \left.\frac{dD}{dE}\right|_{max} \times \Delta E \tag{SI 7}$$

Given the calibration of Grx1-roGFP2 described above equation SI 7 lead to variation in ratio of the order of $|\Delta R_{pH}| \sim 10^{-6}$. Such a variation would affect our readout for adaptation, $\alpha_{add}$, by reducing it $\Delta\alpha \sim 10^{-5}$ which is far smaller than our experimental precision. We concluded that this effect can be fairly neglected in our experimental conditions.

### Single cell parameters to caracterize adaptation dynamics of Grx1-roGFP2 during stimulation

Here we want to characterize cell-to-cell variability in single cell dynamics in response to $100\mu M\ H_2O_2$ step stimulation. For this we fitted each single cell time series and extract parameters. We used a simple phenomenological model considering two superposing processes. First a fast increase of Grx1-roGFP2 upon oxidative stress and second a slower decrease of the ratio. Single cell kinetics were well fitted upon stimulus addition by the following equation :

$$R(t) = A_{add} \times e^{-t/\tau_{add}^D} - B_{add} \times e^{-t/\tau_{add}^R} + C_{add} \quad \text{(SI 8)}$$

where $\tau_{add}^D$ is the ratio decay time and $\tau_{add}^R$ is the ratio rise time. $\tau_{add}^D$ is much slower than $\tau_{add}^R$. $A_{add}$, $B_{add}$ and $C_{add}$ are positive coefficients. Before stimulation $R(t = 0) = A_{add} - B_{add} + C_{add}$ is the basal Grx1-roGFP2 signal. $B_{add}$ is the amplitude by which the ratio is increased shortly after stimulation, it is equal to the empirical parameter $\Delta R_{max} = R_{max} - R(t = 0)$. $A_{add}$ is the amount by which the Grx1-roGFP2 ratio is decreased after prolonged stimulation.

The same equation can be used to fit Grx1-roGFP2 kinetics upon stimulus removal. Similar parameters were defined ($\tau_{rmv}^D$, $\tau_{rmv}^R$, $A_{rmv}$, $B_{rvm}$ and $C_{rmv}$). We note that $A_{rmv}$ and $B_{rmv}$ are negative while $C_{rmv}$ is positive. $B_{rmv}$ is the amplitude by which the ratio is reduced shortly after stimulus removal. $A_{rmv}$ is the amount by which the Grx1-roGFP2 ratio is increased longly after stimulus removal.

In addition we defined two other parameters. $A_{cell}$ is the apparent cell area defined by the number of pixels contained in the mask after single cell segmentation. This parameter reflects cell area in contact with external medium and stimulus. $Fluo$ is the GRx1-roGFP2 fluorescence intensity under $482nm$ excitation which reflects Grx1-roGFP2 expression level.

We first used principal component analysis (PCA) [7] to try to reduce system dimensionality. Indeed if there were strong linear correlation between parameters, new quantities could be defined as linear combination of the parameters to reduce the amount of independent quantities that describe the data. PCA was performed on centered and normalized parameters. For this we substracted each parameters by the mean of the dataset and divided it by the standard deviation.

From the 12 parameters defined above PCA provided a new set of 7 parameters that explain 95% of the observed variance. While this was an improvement, this was not sufficient to further simplify data representation. In order to account for ratio and multiplication in combining parameters we have also used PCA on the logarithm of the parameters with similar results.

We finally reported correlation coefficient of all parameters on Figure SI 3.

[8] Bradley Efron. *The jackknife, the bootstrap, and other resampling plans*, volume 38. Siam, 1982.

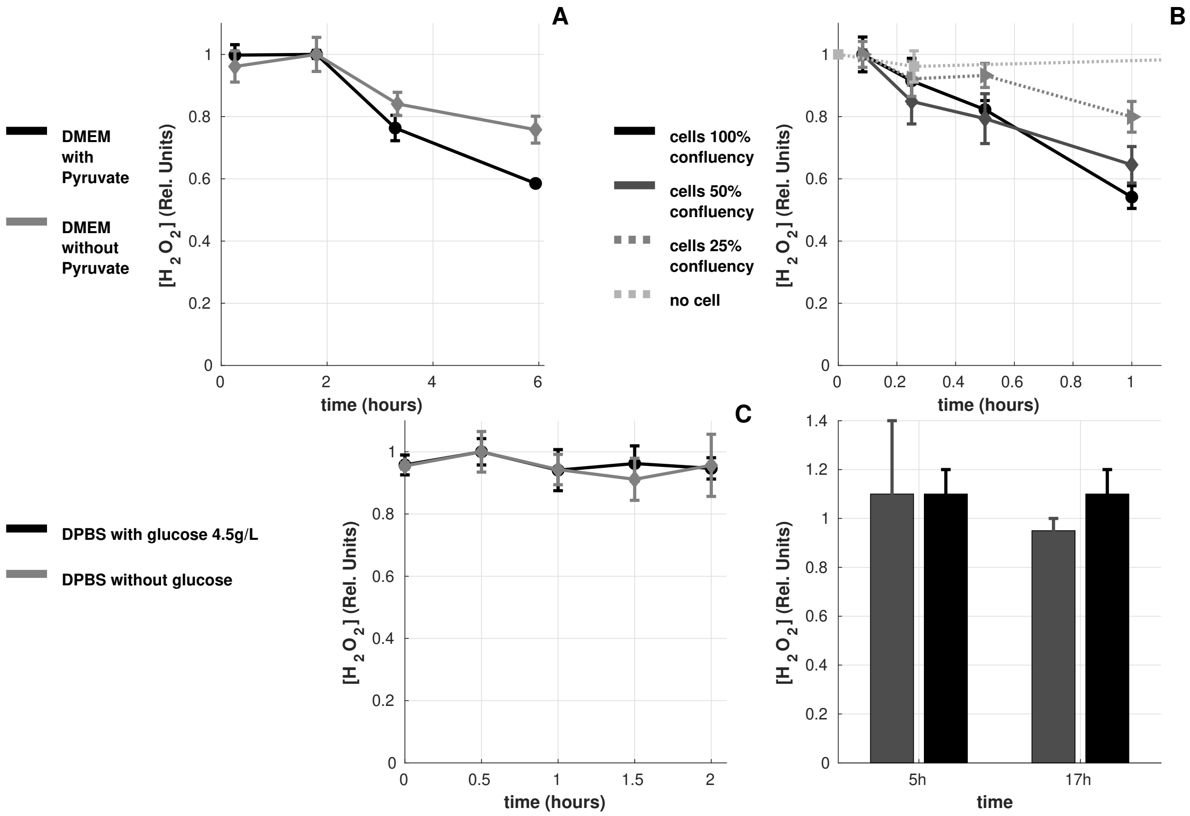


Figure SI 1: **Influence of medium composition on $H_2O_2$ degradation** : We studied kinetics of $H_2O_2$ degradation DMEM with and without Pyruvate, in the presence or absence of cells and in DPBS with or without glucose. We used HyPerBlu to quantify $H_2O_2$ concentration in the various conditions. Data were normalized to maximum HyPerBlu signal. Error bars represent standard deviation. Time evolution of $H_2O_2$ concentration for : **A - complete DMEM** with (black line) and without Pyruvate (gray line) ; **B - cell culture in complete DMEM** with no cells (light gray and dashed line), at 25% confluence (gray and dashed line), 50% confluence (gray and continuous line) and 100% confluence (black and continuous line) ; **C - DPBS** with 4.5g/L glucose (black line) and without glucose (gray line).)

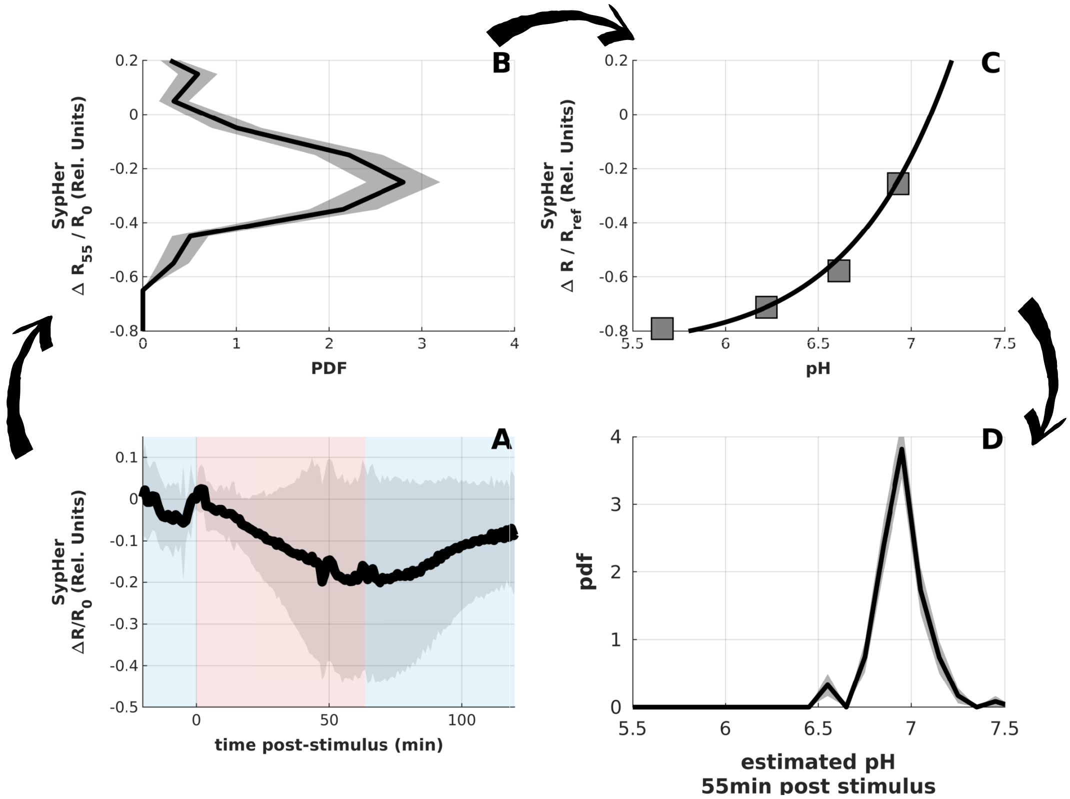


Figure SI 2: **pH estimation upon $H_2O_2$ exposure** : MCF7 cells expressing cytoplasmic SypHer were imaged in the flow chamber. Cells were exposed to $H_2O_2$ free DPBS supplemented with 4.5g/L glucose for 30$min$ then exposed to 100$\mu M$ $H_2O_2$ in DPBS with 4.5g/L glucose for 60$min$ and recovered in $H_2O_2$ free DPBS supplemented with 4.5g/L glucose for another 60$min$. We imaged 166 cells at 1 frame per $min$. **A -** We reported population averaged variation of SypHer ratio, $\Delta R$ relative to pre-stimulus ratio, $R_0$. For single cell data we extracted variation of the ratio 55min post-stimulation, $\Delta R_{55}$. Grey shaded area represents standard deviation. **B -** We report the probability density function of $\Delta R_{55}$ $R_0$. For both PDF from empirical distribution the gray shaded area represents 65% confidence interval obtained by bootstrap resampling [8]. **C -** Calibration of SypHer ratio as a function of pH was extracted from [2]. Data are fitted with equation SI 3 with : $R_{ref} = 0.758$ ; $\Delta R_1 = 0.1$ ; $\Delta R_1 = 9.5$ ; $\nu = 2$ ; $pKa = 8.4$. This calibration allowed us to convert single cell $\Delta R_{55}$ $R_0$ to pH value. **D -** We reported probability density function of estimated pH. Grey shaded area represents 65% confidence interval obtained by bootstrap resampling [8].

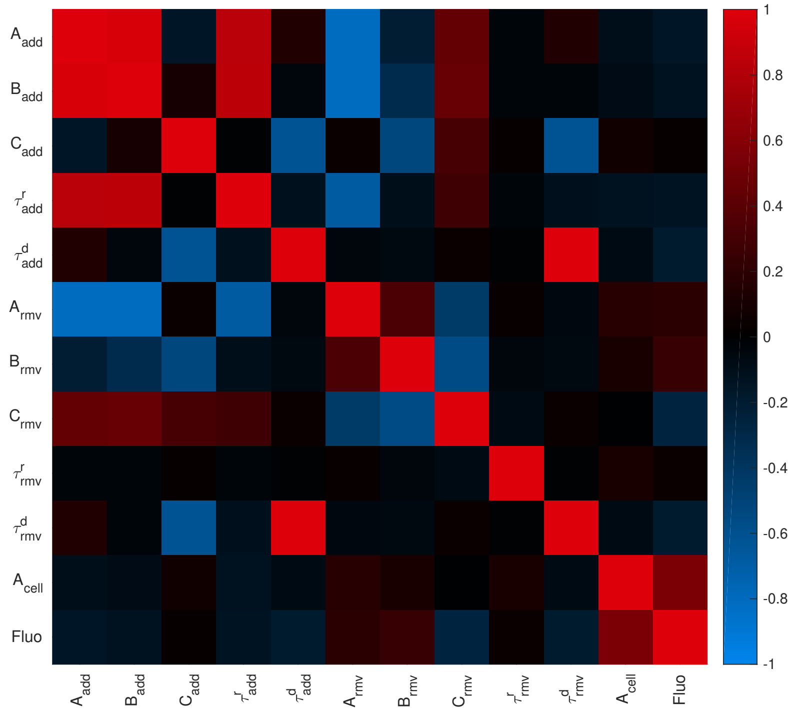


Figure SI 3: Correlation coefficient for fitted parameters. Shape parameters are defined in the text. Correlation coefficient, $r$, of random variables $X$ and $Y$ is defined as : $r = \frac{<XY>-<X><Y>}{\sigma_X \sigma_Y}$ where $< . >$ denotes mean and $\sigma$ standard deviation.

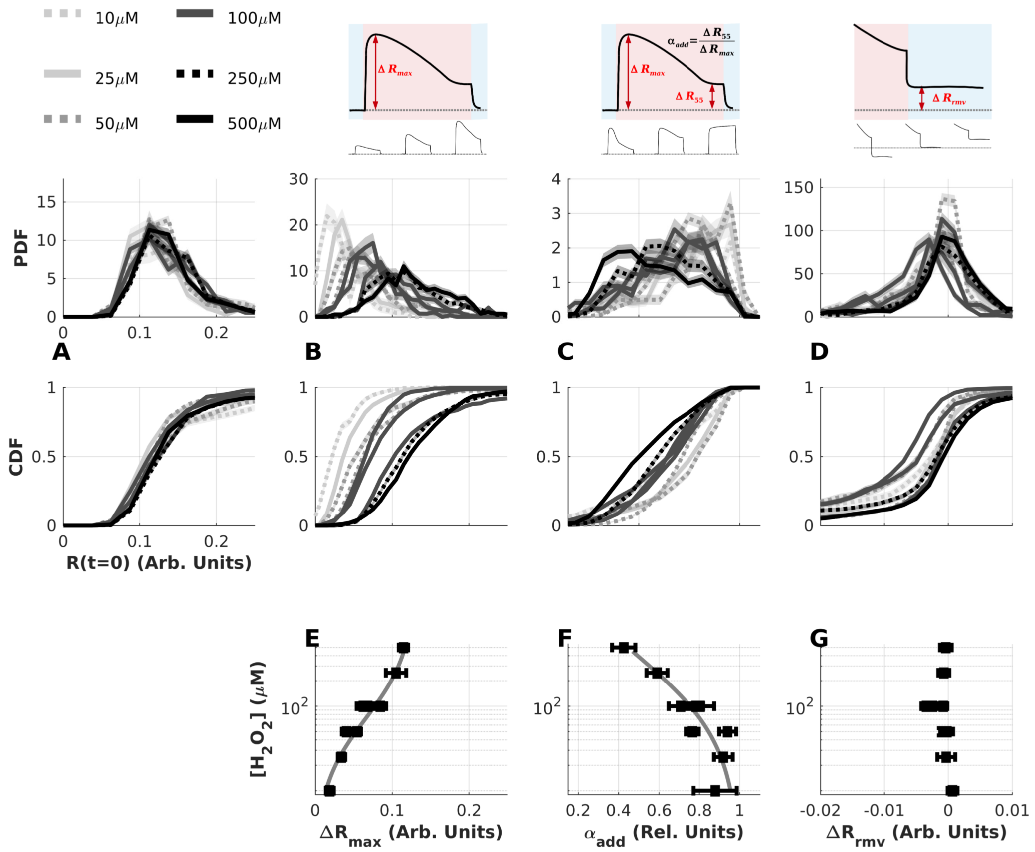


Figure SI 4: **Influence of external $H_2O_2$ concentration on single cell shape parameters** : We repeated experiments reported in Figure 1 at several $H_2O_2$ concentrations ranging from $10\mu M$ to $500\mu M$. In order to quantitatively describe dynamics of $R(t)$ we report shape parameters : **A -** initial state, $R_0$ ; **B -** maximum $\Delta R$ upon stimulus addition ($\Delta R_{max}$) ; **C -** adaptation index ($\alpha_{add}$) defined as the ratio of $\Delta R$ $55min$ after addition and $\Delta R_{add}$ ; **D -** maximum $\Delta R$ upon stimulus removal ($\Delta R_{rmv}$). Upper panels show schematic definition of shape parameters. To account for cell-to-cell variability are reported both the Probability density Function (PDF, middle panel) and the Cumulative Density Function (CDF, lower panel). For CDF the y-axis can be read as the probability to find a cell with a shape parameter higher than x-axis. Data are plotted with gray level representing $H_2O_2$ concentration : lighter gray corresponds to lower concentration and darkness is increasing with $[H_2O_2]$ (see legend). For both PDF and CDF estimated from empirical distribution the gray shaded area represents 65% confidence interval obtained by bootstrap resampling [8]. **E-G - Dose dependence of shape parameters -** We report dependency of the most probable shape parameter as a function of $H_2O_2$ concentration ($[H_2O_2]$). Error bars represent 65% confidence interval obtained by bootstrap resampling [8].
